# Cantharidin alters GPI-anchored protein sorting by targeting Cdc1 mediated remodeling in Endoplasmic Reticulum

**DOI:** 10.1101/460253

**Authors:** Pushpendra Kumar Sahu, Raghuvir Singh Tomar

## Abstract

Cantharidin (CTD) is a potent anticancer small molecule produced by several species of blister beetle. It has been a traditional medicine for the treatment of warts and tumors for many decades. CTD suppresses the tumor growth by inducing apoptosis, cell cycle arrest, and DNA damage. It is a known inhibitor of PP2A and PP1. In this study, we identified new molecular targets of CTD using *Saccharomyces cerevisiae* as a model organism which expresses a Cantharidin Resistance Gene (CRG1). *CRG1* encodes a SAM-dependent methyltransferase that inactivates CTD by methylation. CTD alters lipid homeostasis, cell wall integrity, endocytosis, adhesion, and invasion in yeast cells. We found that CTD specifically affects the phosphatidylethanolamine (PE) associated functions which can be rescued by supplementation of ethanolamine (ETA) in the growth media. CTD also perturbed ER homeostasis and cell wall integrity by altering the GPI-anchored protein sorting. The CTD dependent genetic interaction profile of *CRG1* revealed that Cdc1 activity in GPI-anchor remodeling is the key target of CTD, which we found to be independent of PP2A and PP1. Furthermore, our experiments with human cells suggest that CTD functions through a conserved mechanism in higher eukaryotes as well. Altogether, we conclude that CTD induces cytotoxicity by targeting Cdc1 activity in GPI-anchor remodeling in the endoplasmic reticulum (ER).

## INTRODUCTION

Glycosylphosphatidyl inositol (GPI) anchor biosynthesis is an essential and conserved pathway in eukaryotes. GPI-anchoring is a type of post-translational modification of proteins destined to the plasma membrane or cell wall. This modification takes place in the endoplasmic reticulum (ER) which acts as a signal for sorting of the proteins to the cell surface. The GPI-anchored protein sorting occurs via ER-Golgi traffic system (1). The GPI-anchor biosynthesis takes place in the inner-membrane of ER through a series of enzymatic reactions, which is subsequently incorporated onto the C-terminus of protein (1). Post-synthesis, the GPI-anchor undergoes several steps of modifications in ER (yeast) or Golgi (mammals). This sequential process of modifications is called GPI-anchor remodeling. Bst1, Cdc1, Ted1, Per1, Gup1, and Cwh43 are the key factors, which mediate the process of GPI-anchor remodeling in yeast (1,2). Cdc1 acts as a Mn^+2^-dependent ethanolaminephosphate (EtNP) phosphodiesterase that removes EtNP from the first mannose of the GPI (3,4). *CDC1* is a homolog of human PGAP5 and is essential for cell survival (4,5). Therefore, different point mutants have been created to characterize the *CDC1* function (3,4,6). Previous studies have reported that *cdc1-314* mutant exhibits defect in GPI-anchored protein sorting, temperature sensitivity, cell wall damage, actin depolarization, increased Ca^+2^ ion signaling, and unfolded protein response (UPR) (3,4). GPI-anchored proteins have diverse biological functions in different organisms. In yeast, they regulate cell wall biosynthesis, flocculation, adhesion, and invasion (7). In protozoa (*Trypanosoma brucei)*, GPI-anchored proteins form a protective layer on the cell surface, which helps in the virulence of the parasite (8,9). In plants, it is required for the cell wall biosynthesis, developmental morphogenesis, pollen tube germination, etc. (2). In mammals, it regulates embryogenesis, fertilization, immune response, neurogenesis, etc. (2,9). GPI-anchored proteins are also associated with the progression, invasion, and metastasis of malignant cells (10-12). A few GPI-anchored proteins have been found to serve as markers for the specific stages of tumors (13,14).

Cantharidin (CTD) is a terpenoid produced by blister beetles as a defense molecule. The people of a few Asian countries have been using it as a traditional medicine for the treatment of warts and molluscum contagiosum for more than 2,000 years (15). In the last few decades, many studies have demonstrated the anticancer property of CTD. It has been shown to inhibit the growth of hepatocellular carcinoma (16), leukemia (17), pancreatic (18), colorectal (19), gallbladder (20), oral (21), and breast cancer (22). The serine-threonine protein phosphatases, PP1 and PP2A, are the only reported molecular targets of CTD (23,24). The inhibition of PP2A causes cell cycle arrest (25,26) and apoptosis (27,28). CTD also impairs different cellular processes such as heat shock response (29), autophagy (22), DNA damage response, and mitogen-activated protein kinase (MAPK) signaling (18,21). One of these studies also demonstrated a PP2A or PP1 independent alteration in heat shock response (29), suggesting the existence of additional molecular targets of CTD (29,30). Most of the studies performed with CTD were based on mammalian cell lines, making it difficult to decipher a conserved mechanism of action of the drug due to their tissue-specific origin and differential gene regulation. Hence, yeast (*Saccharomyces cerevisiae*) serves as an appropriate model system to identify the conserved molecular targets of the drug (31-33). Previous studies showed that yeast *YHR209W* gene was essentially required for CTD resistance (34), which was subsequently named as Cantharidin Resistant Gene (*CRG1*) (35). Later on, Crg1 was characterized as an S-adenosylmethionine (SAM) dependent methyltransferase that detoxifies CTD by methylation (30). Deletion of *CRG1* enables the identification of the molecular targets of CTD more easily, so, we utilized budding yeast as a model organism to dissect the molecular mechanism of CTD toxicity.

Our study was focused on the identification of the conserved cellular pathways targeted by CTD. Interestingly, we found that CTD impaired the GPI-anchored protein sorting by targeting the remodeling process in ER. More specifically, it affected the Cdc1 activity leading to multiple cellular changes such as missorting and aggregation of GPI-anchored proteins, temperature sensitivity, cell wall damage, and decreased UPR. Most of the CTD-induced phenotypes observed in yeast cells were also reproducible in human cells. Our comprehensive genetic and cell biology based experiments revealed that the Cdc1 activity is a molecular target of CTD in eukaryotic cells. Overall, we identified the GPI-anchor remodeling as a direct target of CTD.

## RESULTS

### Supplementation of ethanolamine (ETA) suppresses the cytotoxic effect of CTD

Previous studies have shown that CTD treatment affects the lipid homeostasis in budding yeast by inhibition of the elongation of short-chain phospholipids to long-chain phospholipids (30). The phospholipid imbalance can be restored with exogenous supplementation of the precursor molecules. For example, supplementation of ETA and choline (CHO) activates the synthesis of phosphatidylethanolamine (PE) and phosphatidylcholine (PC), respectively, via an alternative pathway, i.e., the Kennedy Pathway (Fig. 1F) (36). Inositol (INO) and serine (SER) enter into the canonical pathways of phosphatidylinositol (PI) and phosphatidylserine (PS) biosynthesis, respectively (Fig. 1F) (37,38). Based on these phenomena, we sought to identify the specific phospholipid affected by CTD. We supplemented the medium with specific precursor molecules, ETA, CHO, and INO, with or without CTD and measured the growth of wild type (WT) and *crg1Δ* strains (Fig. 1A). CTD exposure produced a lethal effect on *crg1Δ* mutant compared to WT (30). However, ETA supplementation completely rescued the growth of the *crg1Δ* strain from CTD cytotoxicity (Fig. 1A, S1A–D). On the other hand, CHO and INO supplementation failed to rescue the growth of *crg1Δ* strain in CTD-containing medium (Fig. 1A). This observation suggests that CTD specifically targets PE. The exclusive rescue in the growth of the CTD-treated cells by ETA supplementation was a surprising phenomenon, because PE and PC, both are synthesized in the same pathway (37). Thus, we believe that CTD may not affect the PE biosynthesis pathway, but it might be altering the PE-associated structures or functions. PE plays an essential role in maintaining membrane and cell wall integrity under heat stress (38,39); so, we examined the fitness profile of WT and *crg1Δ* strains in heat stress (37°C) with a permissible dose of CTD (2μM). Interestingly, we found complete inhibition of growth of *crg1Δ* mutant at 37°C in the presence of CTD, whereas the growth was unaffected at optimum (30°C) or below the optimum (25°C) temperature (Fig. 1B). CTD cytotoxicity was suppressed again at 37°C by supplementation of ETA (Fig. 1C). PE biosynthesis takes place in mitochondria and Golgi/vacuole with the help of Psd1 and Psd2, respectively (40). A major fraction (>90%) of the net PE in a cell is synthesized by Psd1 in mitochondria (40), so we created a double-deletion mutant, *crg1Δpsd1Δ,* to check synthetic lethality between *PSD1* and *CRG1* in the presence of CTD. For this purpose, WT, *crg1Δ, psd1Δ*, and *crg1Δpsd1Δ* strains were grown in CTD-containing medium. We found that the *crg1Δpsd1Δ* mutant was hypersensitive to CTD than *crg1Δ*, suggesting that PE is essentially required to tolerate CTD toxicity (Fig. 1D, S1E–H). *crg1Δpsd1Δ* mutant followed the same trend at higher temperature as well (37°C) (Fig. 1E). The synthetic lethality between *CRG1* and *PSD1* in the presence of CTD suggests an essential role of PE to tolerate CTD toxicity. These observations suggest that CTD affects the PE-associated functions (Fig. 1F); therefore, enhanced synthesis of PE helps to overcome the CTD toxicity.

**Figure 1:**
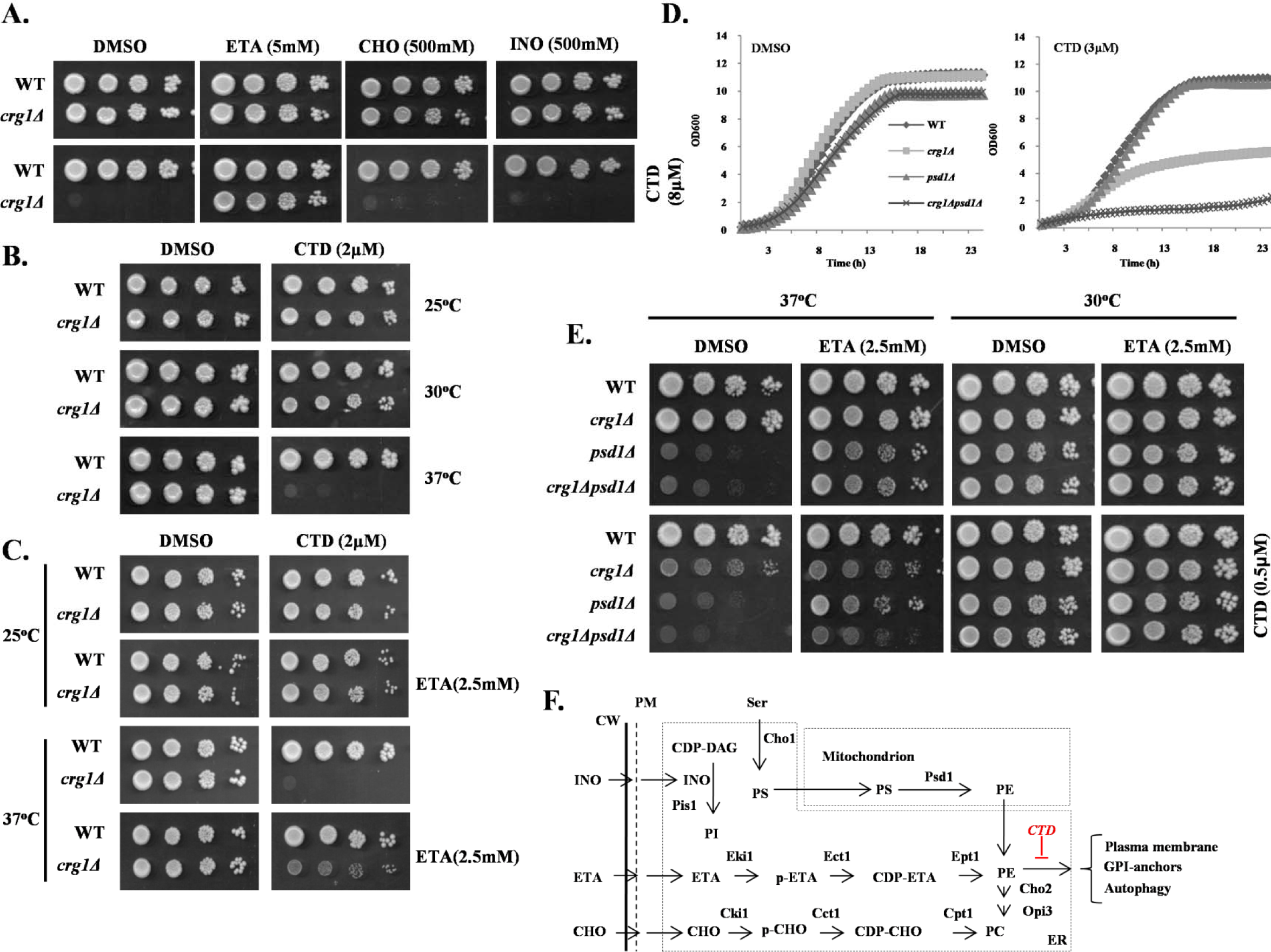
CTD specifically targets PE in *crg1Δ* cells. A, B, C, and E: Growth sensitivity assays. Equal number of cells were serially diluted and spotted on SC agar media. Images were captured after 72h of incubation. (A) Supplementation of ETA rescues *crg1Δ* mutant from CTD toxicity. Phospholipid precursors: ETA, INO, and CHO were added into SC agar media with or without CTD. WT and *crg1Δ* cells spotted and incubated at 30°C. (B) CTD toxicity increases with rising temperature .WT and *crg1Δ* cells were spotted on SC agar media containing CTD and incubated at different temperatures (25°C, 30°C, 37°C). (C) ETA supplementation rescues the *crg1Δ* mutant from CTD toxicity at higher temperature. WT and *crg1Δ* cells were spotted on SC agar media containing CTD with and without ETA supplementation, and incubated at different temperatures (25°C, 30°C, 37°C). (D&E) *CRG1* shows synthetic lethality with *PSD1* under CTD stress. (D) Growth curve assay. Equal number of cells of WT, *crg1Δ, psd1Δ* and *crg1Δpsd1Δ* were grown at 30°C with or without CTD in liquid media. OD_600_ was measured at the time interval of 30 minutes using an automated plate reader for 23h. (E) WT, *crg1Δ, psd1Δ* and *crg1Δpsd1Δ* cells were spotted on SC agar media containing CTD with or without ETA, and incubated at two different temperatures (25°C and 37°C). (F) Phospholipid biosynthesis pathways in yeast (37,66,77,78). INO and SER in media are directly utilized to synthesize PI and PS with help of Pis1 and Cho1, respectively. PE and PC biosynthesis has two pathways: (i) Canonical biosynthesis of PE/PC: It takes place in mitochondria and ER. The first reaction starts in ER where Cho2 synthesizes PS from serine (SER). PS is transported to mitochondria where Psd1 catalyses its decarboxylation to synthesize PE. (Similar mechanism also takes place in Golgi and vacuole by Psd2 which contributes very minor fraction of the net PE content). Next, PE is transported again to ER where Cho2 and Opi3 convert it into PC via a sequence of methylation reactions. (ii) Non-canonical PE or PC synthesis: It is also called as Kennedy pathway. In this pathway, externally supplemented precursors (ETA/CHO) are utilized and converted into PE or PC, respectively, via series of enzymatic reactions.

### CTD alters ER homeostasis by inhibition of UPR

ER is the organelle for the synthesis of the major phospholipids. Imbalance in the phospholipid composition of lipid bilayer membrane is reported to induce ER stress (41-44). Existing evidence suggests that CTD also perturbs ER-synthesized phospholipids (30); thus, we proposed that CTD might be altering the ER homeostasis. We examined ER stress in *crg1Δ* cells in the presence of CTD. Firstly, WT and *crg1Δ* cells were co-treated with CTD and ER stress (or UPR) inducers, dithiothreitol (DTT) or tunicamycin (TM), to check if there was any synergistic effect between the two molecules. For this, we chose a permissible dose of CTD (4μM) for the *crg1Δ* mutant, at which it survived, but survival was lower than that of the WT. Both the strains were spotted on CTD-containing medium, with and without TM or DTT. Interestingly, the co-treatments (CTD + TM and CTD + DTT) inhibited the growth of *crg1Δ* cells more severely compared to only CTD treatment (Fig. 2A, S1I–L). The synergistic lethal effect on the growth of *crg1Δ* cells upon co-treatments suggests that CTD perturbs ER homeostasis. Next, we measured UPR by β-galactosidase assay with the help of UPRE-LacZ reporter plasmid (45,46). We found that the basal level of UPR was lower in *crg1Δ* cells compared to WT, and CTD treatment further inhibited UPR in both the strains (WT and *crg1Δ*) (Fig. 2B). Since ETA supplementation rescues the yeast cells from CTD toxicity, we decided to measure UPR upon ETA supplementation. Surprisingly, ETA supplementation could not rescue UPR inhibited by CTD (Fig. 2B), suggesting that CTD inhibits UPR via a distinct mechanism independent of PE in ER. To get more insight into this mechanism, UPR was measured upon co-treatments of cells with CTD+DTT and CTD+TM. We found decreased UPR levels in *crg1Δ* cells upon DTT and TM treatment (Fig. 2C). Moreover, CTD treatment strongly inhibited UPR induced by DTT or TM in WT as well as *crg1Δ* mutant (Fig. 2C). Consistent with these observations, we also found a slight inhibition of *HAC1* mRNA splicing in *crg1Δ* cells compared to WT. The splicing of *HAC1* mRNA was further inhibited in both the strains, WT and *crg1Δ*, upon CTD treatment (Fig. 2D). DTT and TM treatment strongly induced *HAC1* mRNA splicing; however, the presence of CTD with DTT or TM suppressed *HAC1* mRNA splicing strongly in *crg1Δ* cells (Fig. 2D). These results suggest that CTD inhibits UPR by making an obstruction in *HAC1* mRNA splicing, although the mechanism remains unclear.

**Figure 2:**
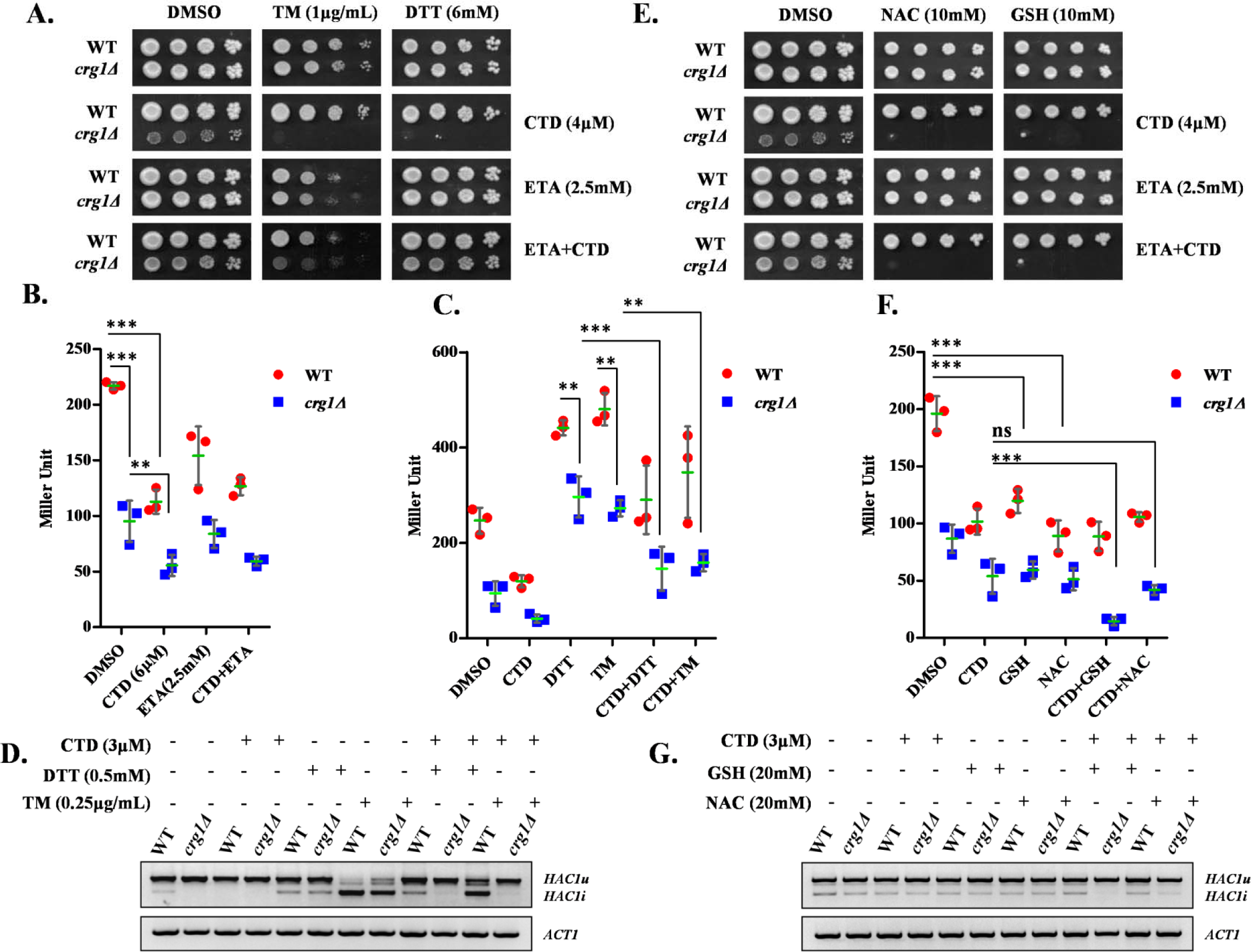
CTD treatment inhibits UPR by alteration of the ER-redox homeostasis. (A) UPR inducers (DTT/TM) synergistically enhance CTD toxicity. Equal number of serially diluted WT and *crg1Δ* cells were spotted on CTD containing SC agar media with or without DTT/TM in presence or absence of ETA, and incubated at 30°C for 72h. (B) CTD inhibts UPR. WT and *crg1Δ* strains transformed with pPW344 (UPRE-LacZ) plasmid, were grown in SC-URA media at 30°C. Cells were treated with CTD (6μM) with or without ETA (2.5mM) at the mid-exponential phase (OD_600_ = 0.8) and incubated for 2h. β-galactosidase assay was performed to measure the UPR. The graph shows scattered plot of each data point of individual experiment (dot/square) with mean (horizontal green line) ± SD (error bars). Statistical analysis was done with GraphPad Prism-5, applying Two-way ANOVA, Bonferroni post test where *= p<0.05, **= p<0.01 and ***= p<0.001 and ns=p>0.05. (C) CTD inhibits UPR in presence of DTT and TM. WT and *crg1Δ* strains carrying pPW344 vector were grown till the mid-exponential phase and treated with CTD (3μM) in combination of DTT (0.5mM) or TM (0.25μg/mL) for 2h. β-galactosidase assay was performed to measure the UPR. The graph shows scattered plot of each data point of individual experiment (dots/squares) with mean (green line) ± SD (error bars). Statistical analysis was done using GraphPad Prism-5, applying Two-way ANOVA, Bonferroni post test where *= p<0.05, **= p<0.01 and ***= p<0.001 and ns=p>0.05. (D) CTD inhibits *HAC1* mRNA splicing. WT and *crg1Δ* strains were grown in the condition mentioned above (C) and *HAC1* mRNA splicing was measured by RT-PCR. (E) GSH or NAC supplementation enhances the CTD cytotoxicity. Equal number of WT and *crg1Δ* cells were serially diluted and spotted on CTD containing SC agar media with or without reducing agents (GSH and NAC) in presence or absence of ETA, incubated at 30°C for 72h. (F) GSH and NAC supplementation reduces UPR. WT and *crg1Δ* strains transformed with pPW344 (*UPRE-LacZ)* were grown in SC-URA media at 30°C till mid-exponential phase. The cells were treated with CTD (3μM) in presence of absence of GSH (20mM) or NAC (20mM) for 2h and processed for β-galactosidase assay. The graph shows scattered plot of each data point of individual experiment with mean±SD (error bars). Statistical analysis was done using GraphPad Prism-5, applying Two-way ANOVA, Bonferroni post test where *= p<0.05, **= p<0.01 and ***= p<0.001 and ns=p>0.05. (G) GSH and NAC supplementation enhances the CTD mediated inhibition of *HAC1* splicing. WT and *crg1Δ* cells were grown in SC media at 30°C till mid-exponential phase under same conditions mentioned above (F) and the *HAC1* mRNA splicing was measured by RT-PCR.

Our results suggest that CTD exposure leads to ER stress that cannot be rescued by ETA supplementation. The ER-lumen maintains higher oxidation potential with the help of low GSH:GSSG ratio (1:1 to 3:1) compared to the high GSH:GSSG ratio (30:1 to 100:1) of the cytosol (47). GSH provides a redox buffer for the catalytic activity of the protein-folding enzymes in ER (48,49). The imbalance in GSH:GSSG ratio in ER impairs oxidative protein folding that causes ER stress (50,51). Based on these previous findings, we predicted that CTD induced ER stress might be due to imbalance in GSH:GSSG ratio in ER. To test this hypothesis, we checked the effect of GSH on CTD toxicity. We used the permissible dose of CTD (4μM) for *crg1Δ* mutant and supplemented the medium with a high dose of GSH and NAC. We found that the growth of *crg1Δ* mutant was suppressed in the presence of either of the two reducing molecule, GSH or NAC, along with CTD. However, GSH or NAC alone did not show any effect on the growth of *crg1Δ* mutant (Fig. 2E). Furthermore, the supplementation of ETA did not rescue the growth of *crg1Δ* mutant upon CTD+GSH or CTD+NAC treatments. This result supports the previous observation where, upon ETA supplementation, UPR suppressed by CTD treatment could not be rescued (Fig. 2B). Similar observations were also made in liquid growth culture (Fig. S2A–F). Next, we measured UPR using β-galactosidase assay. Interestingly, we observed that GSH or NAC supplementation results in the reduction in UPR in WT and *crg1Δ* cells (Fig. 2F). As CTD treatment also inhibits UPR, we observed severe reduction in UPR upon co-treatments with CTD+GSH or CTD+NAC (Fig. 2F). We also found an inhibition in *HAC1* mRNA splicing upon addition of GSH and NAC (Fig. 2G). The splicing of *HAC1* mRNA decreased robustly when the cells were co-treated with CTD+GSH or CTD+NAC (Fig.2G). These observations suggest that CTD-mediated inhibition of UPR is probably due to imbalance in ER-redox homeostasis, which gets enhanced with the addition of GSH. It also explains the reason why ETA supplementation failed to rescue the UPR.

### CTD exposure perturbs the cell wall integrity via ER stress

Yeast cell wall biosynthesis and maintenance largely depend on functional ER (7,52,53). Dysfunctional ER affects the synthesis, modifications, folding, and transport of the proteins destined to the plasma membrane or cell wall. Based on our results, we proposed that CTD induced ER stress could also perturb cell wall integrity. To examine the effects of CTD on cell wall integrity, we measured chitin content in the cell wall of WT and *crg1Δ* cells by Calcofluor white (CFW) staining (54). We found substantial increase in chitin content in *crg1Δ* cells upon CTD treatment, suggesting that CTD treatment induced cell wall damage (Fig. S3A). To get more insight on the effect of CTD on cell wall integrity, we co-treated the cells with CTD and cell wall perturbing agents, Congo red (CR) and CFW. We used a permissible dose of CTD (4μM) in combination with cell wall perturbing agents to measure the growth of WT and *crg1Δ* cells. We found that *crg1Δ* mutant did not grow in either of the co-treatments (CTD+CR or CTD+CFW), whereas the growth was unaffected in individual treatments (Fig. 3A). We also supplemented sorbitol (SRB) into the medium to maintain the osmotic balance across the cell membrane. SRB rescued the growth of *crg1Δ* mutant upon CTD+CFW treatment, but not upon CTD+CR treatment. That suggests the CTD+CR-induced cell wall damage is irreversible, although the mechanism remains to be identified (Fig. 3A). We obtained similar results in liquid growth culture under similar conditions (Fig. S3B–E). Yeast cell wall damage is sensed by many sensor proteins residing in the cell wall, which in turn activate downstream signaling via Slt2 (55). Activation of Slt2 triggers the transcription of cell wall maintenance genes via Rlm1 and Swi4-Swi6 transcription factors (52,56). Hence, we did western blot analysis of Slt2 phosphorylation in WT and *crg1Δ* cells upon CTD treatment. We observed increased phosphorylation of Slt2 in *crg1Δ* cells upon CTD treatment at 25°C. Slt2 phosphorylation increased further when the cells were grown at 37°C, and CTD treatment induced Slt2 phosphorylation robustly in *crg1Δ* cells at this temperature (Fig. 3B). As we knew that CTD cytotoxicity could be neutralized by ETA supplementation, we decided to measure Slt2 phosphorylation in CTD-treated cells supplemented with ETA. We observed a significant decrease in Slt2 phosphorylation upon ETA supplementation in CTD-treated cells (Fig. 3B). This result suggests that ETA-mediated rescue in growth against CTD toxicity (Fig. 1A, C and Fig. S1A–D) might occur via cell wall maintenance. Next, we challenged the WT and *crg1Δ* cells with the combined treatment of CTD and UPR inducers (DTT and TM) to measure the synergistic effect on Slt2 phosphorylation. We found strong induction in Slt2 phosphorylation upon co-treatments with CTD+DTT or CTD+TM compared to individual treatments (CTD/DTT/TM) (Fig. 3C). This observation suggests that CTD-induced cell wall damage might be due to ER stress. Furthermore, we checked Slt2 phosphorylation upon co-treatments with CTD+GSH and CTD+NAC. We found that both the co-treatments did not cause any significant change in Slt2 phosphorylation compared to CTD alone. Moreover, only GSH or NAC did not cause any change in Slt2 phosphorylation (Fig. 3D), suggesting some unknown mechanism of GSH-induced ER stress unlike DTT, TM, and CTD. We conclude that CTD perturbs cell wall integrity via ER stress.

**Figure 3:**
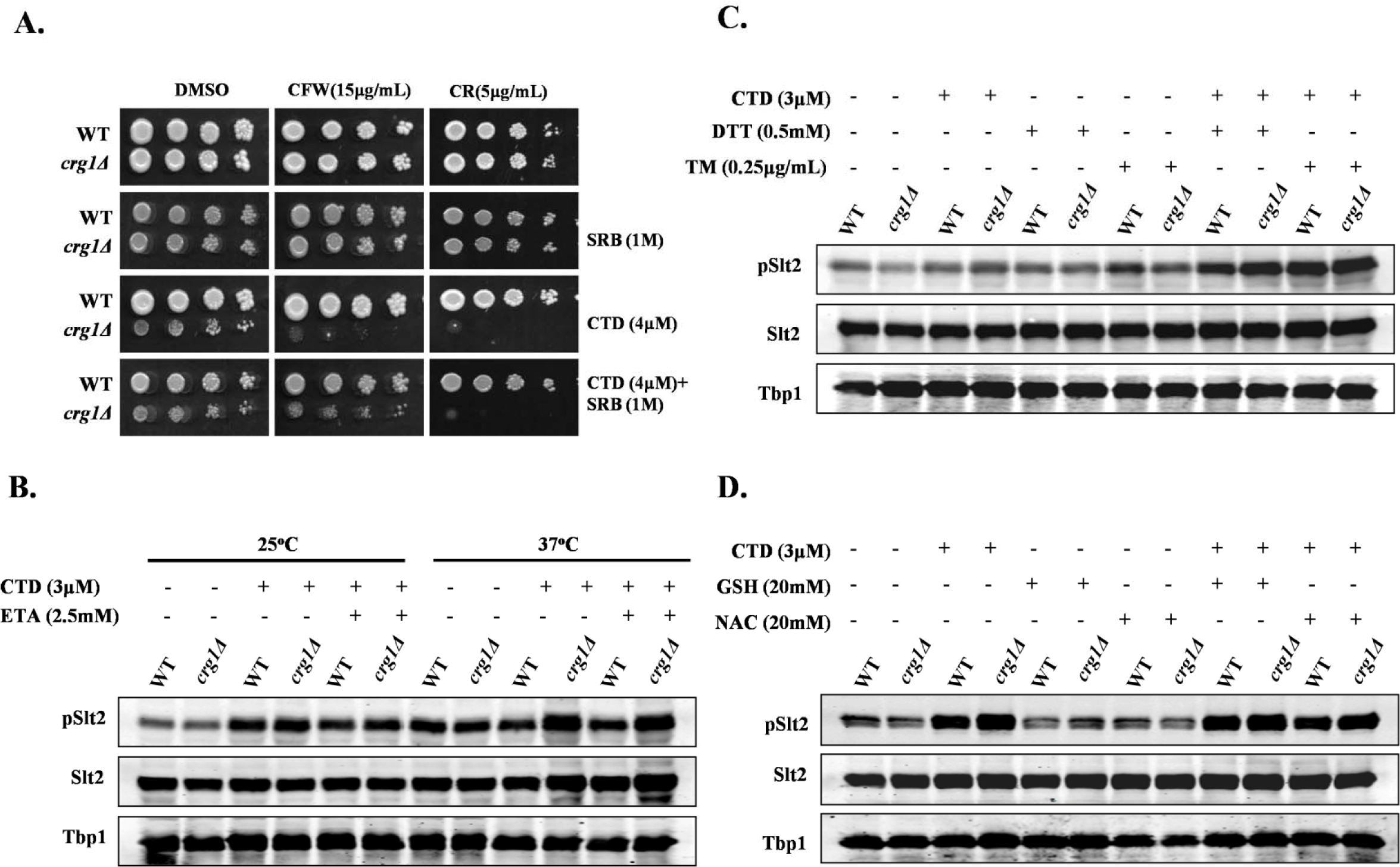
CTD induced ER-stress perturbs the cell wall integrity. (A) CTD and cell wall perturbing agents (CR or CFW) are synergistically lethal to *crg1Δ* mutant. Equal number of WT and *crg1Δ* cells were serially diluted and spotted on SC agar media containing CTD with and without CR or CFW. The cells were incubated at 30°C for 72 h. B, C, and D: Western blot analysis of the Slt2 phosphorylation. Whole cell lysates were prepared from WT and *crg1Δ* cells grown in different conditions. Tbp1 was taken as a loading control. (B) CTD induced cell wall damage increases with heat stress. WT an *d crg1Δ* strains were grown at two different temperatures, 24°C and 37°C till mid-exponential phase (0.8 OD_600_) then treated with CTD in presence or absence of ETA for 2h. (C) CTD induced cell wall damage increases with UPR induction. WT and *crg1Δ* strains were grown at 24°C till mid-exponential phase and treated with CTD with or without DTT or TM for 2h. (D) GSH or NAC supplementation doesn’t affect CWI pathway. WT and *crg1Δ* strains were grown at 24°C till mid-exponential phase, and then treated with CTD in presence or absence of GSH and NAC for 2h.

### CTD alters GPI-anchored protein sorting

To identify the major pathway affected by CTD treatment, we did functional clustering (57) of the genetic interactors of *CRG1* which show synthetic lethality in the presence of CTD (30). We found that the majorly affected pathways were associated with ER-Golgi traffic system (Table S4). Yeast cell wall biosynthesis and maintenance mainly depend on the GPI-anchored proteins, sorted by ER-Golgi traffic system (1,7,58). PE also plays a crucial role in the regulation of this traffic system (4,7,44,58). Thus, we hypothesized that the CTD-induced cell wall damage might be due to the defect in GPI-anchored protein sorting. We decided to study the GPI-anchored protein sorting upon CTD treatment. We used Gas1-GFP as a model GPI-anchored protein and tracked its localization in WT and *crg1Δ* cells upon CTD treatment (3,4,59). We observed that CTD induced missorting and aggregation of Gas1-GFP in *crg1Δ* cells (Fig. 4A). Additionally, Gas1-GFP expression decreased significantly after CTD treatment in *crg1Δ* mutant probably due to the degradation of the aggregated proteins (Fig. 4B) (60-62). We also observed band shift of Gas1-GFP upon CTD treatment in *crg1Δ* mutant, suggesting a distinct mechanism affecting the maturity of GPI-anchored proteins (Fig. 4B). Supplementation of ETA completely rescued the sorting of Gas1-GFP in CTD-treated *crg1Δ* cells (Fig. 4A). This might be the reason for ETA-mediated rescue in cell wall integrity (Fig. 3B). Furthermore, we measured DTT extractable cell surface proteins (CSPs) integrated into the cell wall and plasma membrane through GPI-anchors. These proteins were extracted as described previously(63). We found a high yield of CSPs from CTD-treated *crg1Δ* cells compared to WT cells. However, the supplementation of ETA restored the cell surface proteins to normal level, equal to that of WT (Fig. S4A, B). High yield of CSPs from CTD-treated *crg1Δ* cells maybe due to inappropriate anchorage to cell wall or cell membrane, hence, they are easily extractable from the surface. In contrast, ETA supplementation re-stabilizes the binding of the GPI-anchors, reversing the phenotype to normal and equivalent to WT. To further ascertain the role of CTD on GPI-anchor biosynthesis, we checked the genetic interaction of *CRG1* with a few GPI anchor biosynthesis genes (*GPI2, GPI13,* and *MCD4*) (1). Since, these genes are essential for the cell survival, we used their heterozygous deletion mutants (*gpi2Δ/GPI2, gpi13Δ/GPI13*, and *mcd4Δ/MCD4*). We deleted *CRG1* to create double-deletion mutants, *crg1Δ/Δgpi2Δ/GPI2, crg1Δ/Δgpi13Δ/GPI13*, and *crg1Δ/Δmcd4Δ/MCD4*, and performed growth assay uponCTD treatment. Surprisingly, the double-deletion mutants (*crg1Δ/Δgpi2Δ/GPI2, crg1Δ/Δgpi13Δ/GPI13*, and *crg1Δ/Δmcd4Δ/MCD4*) showed better growth compared to single-deletion mutant, *crg1Δ/Δ*, in CTD-treated medium (Fig. 4C, D and Fig. S3F–I). That suggests that the molecular target of CTD maybe downstream of the GPI biosynthesis cascade (4). We conclude that CTD alters the GPI-anchored protein sorting which can be rescued by ETA supplementation.

**Figure 4:**
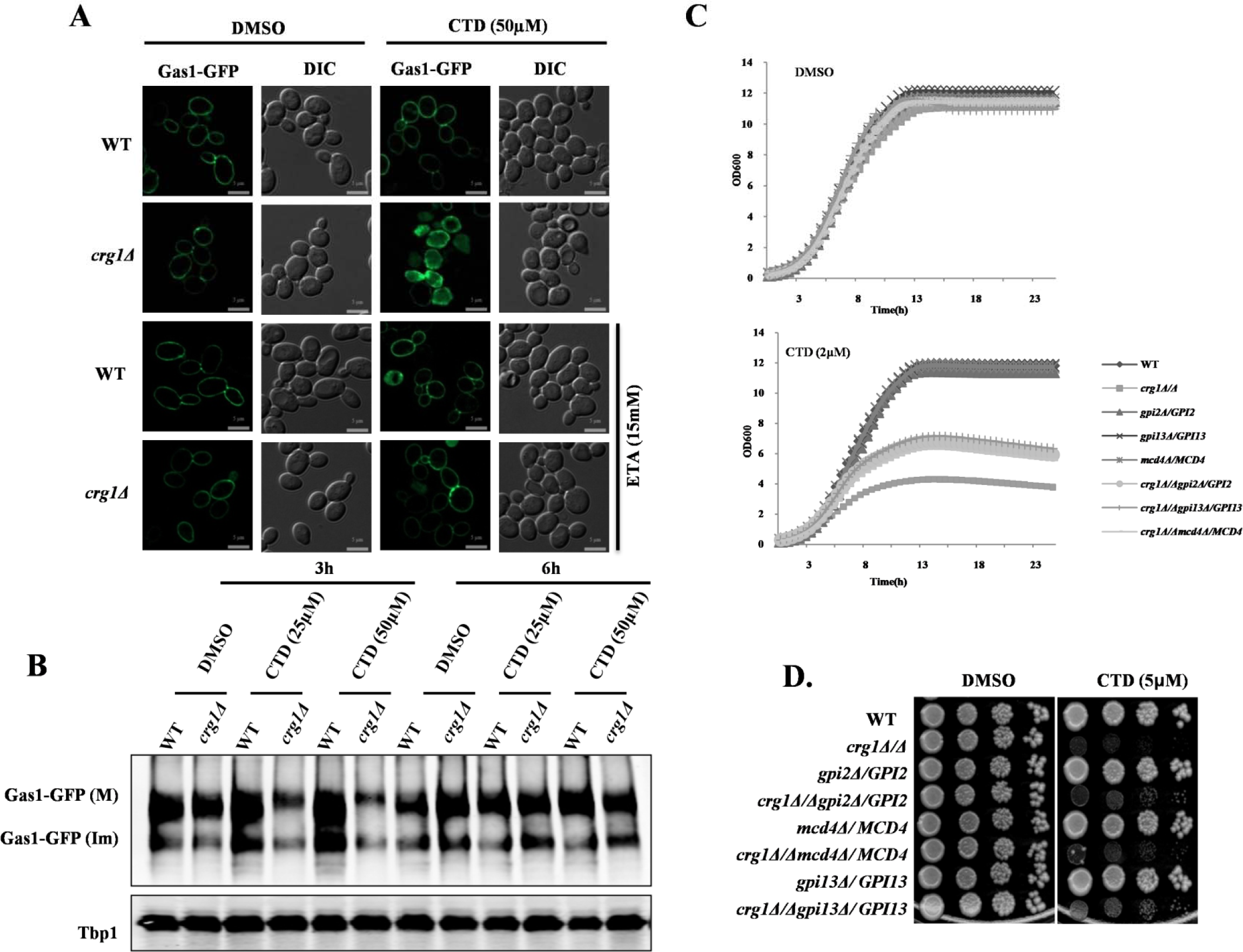
CTD alters GPI-anchored protein sorting. (A) CTD treatment induces missorting of Gas1-GFP. WT and *crg1Δ* cells were transformed with YEp24-GAS1-GFP plasmid. Cells were grown in YPD at 30°C till mid exponential phase, treated with CTD with or without ETA and incubated for 6h before imaging. Sub-cellular localization of Gas1-GFP was observed by using ZEISS-Apotome fluorescence Microscope. (B) CTD treatment decreases Gas1-GFP expression. WT and *crg1Δ* strains expressing Gas1-GFP were grown in YPD at 30°C till mid exponential phase, and then treated with CTD. Cells were harvested after 3h and 6h incubation to analyze the expression of Gas1-GFP using anti-GFP antibody. Tbp1 was used as a loading control. (C&D) GPI biosynthesis genes show synthetic rescue with *CRG1* under CTD stress. (C) Growth curve assay to compare the sensitivity of *crg1Δ/Δ, gpi2Δ/GPI2, gpi13Δ/GPI13, mcd4Δ/MCD4, crg1Δ/Δgpi2Δ/GPI2, crg1Δ/Δgpi13Δ/GPI13* and *crg1Δ/Δmcd4Δ/MCD4* mutants to CTD. (D) Growth sensitivity spot assay of WT, *crg1Δ/Δ, gpi2Δ/GPI2, gpi13Δ/GPI13, mcd4Δ/MCD4, crg1Δ/Δgpi2Δ/GPI2, crg1Δ/Δgpi13Δ/GPI13* and *crg1Δ/Δmcd4Δ/MCD4* mutants. Equal number of WT and mutant cells were serially diluted and spotted on the CTD containing SC agar media. The spotted cells were incubated at 30°C for 72 hr.

### Cdc1-mediated GPI-anchor remodeling is the major target of CTD

GPI-anchor remodeling is the step successive to biosynthesis. Based on the results discussed above, we hypothesized that GPI-anchor remodeling might be the target of CTD (30). We performed experiments to find synthetic lethality between *CRG1* and GPI-anchor remodeling genes (*CDC1, PER1*, and *GUP1*). Firstly, we deleted *CRG1* in the mutants of remodeling factors (*per1Δ, gup1Δ, cdc1-310, cdc1-314, per1Δcdc1-314,* and *gup1Δcdc1-314*) and measured their fitness profile upon CTD treatment. We found that the double mutants (*crg1Δcdc1-314, crg1Δcdc1-310, crg1Δgup1Δ,* and *crg1Δper1Δ*) were hypersensitive to CTD compared to the single mutants (*crg1Δ, per1Δ, gup1Δ, cdc1-310,* and *cdc1-314*) (Fig. 5A). Additionally, the triple mutants (*crg1Δper1Δcdc1-314* and *crg1Δgup1Δcdc1-314*) showed even more sensitivity to CTD than single or double mutants (Fig. 5A and Fig. S5). Interestingly, two different alleles of *CDC1* (*cdc1-310* and *cdc1-314*) showed contrasting phenotypes.*cdc1-314* showed synthetic lethality,whereas *cdc1-310* showed a dose-dependent behavior. It showed synthetic rescue at lower dose and synthetic lethality at higher dose (Fig. 5A and Fig. S5). We also checked their fitness profile upon CTD treatment at a higher temperature (37°C), and we found that *cdc1* mutants showed temperature sensitivity, while the growth of *per1Δ* and *gup1Δ* was unaffected (Fig. 5B) (3,4). However, the double mutants *crg1Δgup1Δ* and *crg1Δper1Δ* were found to be hypersensitive compared to the single-deletion mutant *crg1Δ* at a very low dose of CTD (0.25μM) (Fig. 5B). The results suggest that *CRG1* shows synthetic lethality with GPI-anchor remodeling genes, stronger with *CDC1* than *PER1* or *GUP1* in the presence of CTD. Based on our results and genetic interaction study performed between *CDC1* and *MCD4* (4), we propose that down regulation of GPI-anchor biosynthesis genes or decreased GPI biosynthesis can rescue the growth defect of the mutants lacking GPI-anchor remodeling, perhaps by decreasing the GPI traffic on the remodeling factors. Because of the dynamic behavior of *cdc1* alleles (*cdc1-314* and *cdc1-310*) against CTD, we hypothesized that the Cdc1 activity could be a specific target of CTD in the remodeling process (Fig. 5A and Fig. S5, S6C–F). As the activity of Cdc1 is Mn^+2-^dependent (3,5,6), we decided to examine the effect of CTD by controlling Mn^+2^ concentration in the medium. We added the di-ionic chelator EGTA into the medium along with CTD and checked the fitness profile of the mutants. We observed that the growth of single and double mutants (*crg1Δ, crg1Δcdc1-310*, and *crg1Δcdc1-314*) was suppressed gradually with increasing concentration of EGTA (Fig. 5C and Fig. S6A). Moreover, exogenous supplementation of MnCl_2_recovered the growth of *crg1Δcdc1-310* mutant in CTD-treated medium (Fig. 5D and Fig. S6B, G–J). In this study, the two *cdc1* alleles exhibit reproducible phenotype in a distinct condition than that reported previously (3,6). In summary, we conclude that *CRG1* and *CDC1* work in two different axes: *CRG1* works as a guard to resist CTD, while *CDC1* participates in the remodeling process. Loss of *CRG1* results in the increased availability of active CTD that impaired the remodeling process by targeting the Cdc1 activity (Fig. 5E). Although both the genes work in two different axes, they are required in parallel to tolerate CTD toxicity. We conclude that Cdc1 activity is essential to tolerate CTD cytotoxicity, and it may serve as a mechanistic target of the drug.

**Figure 5:**
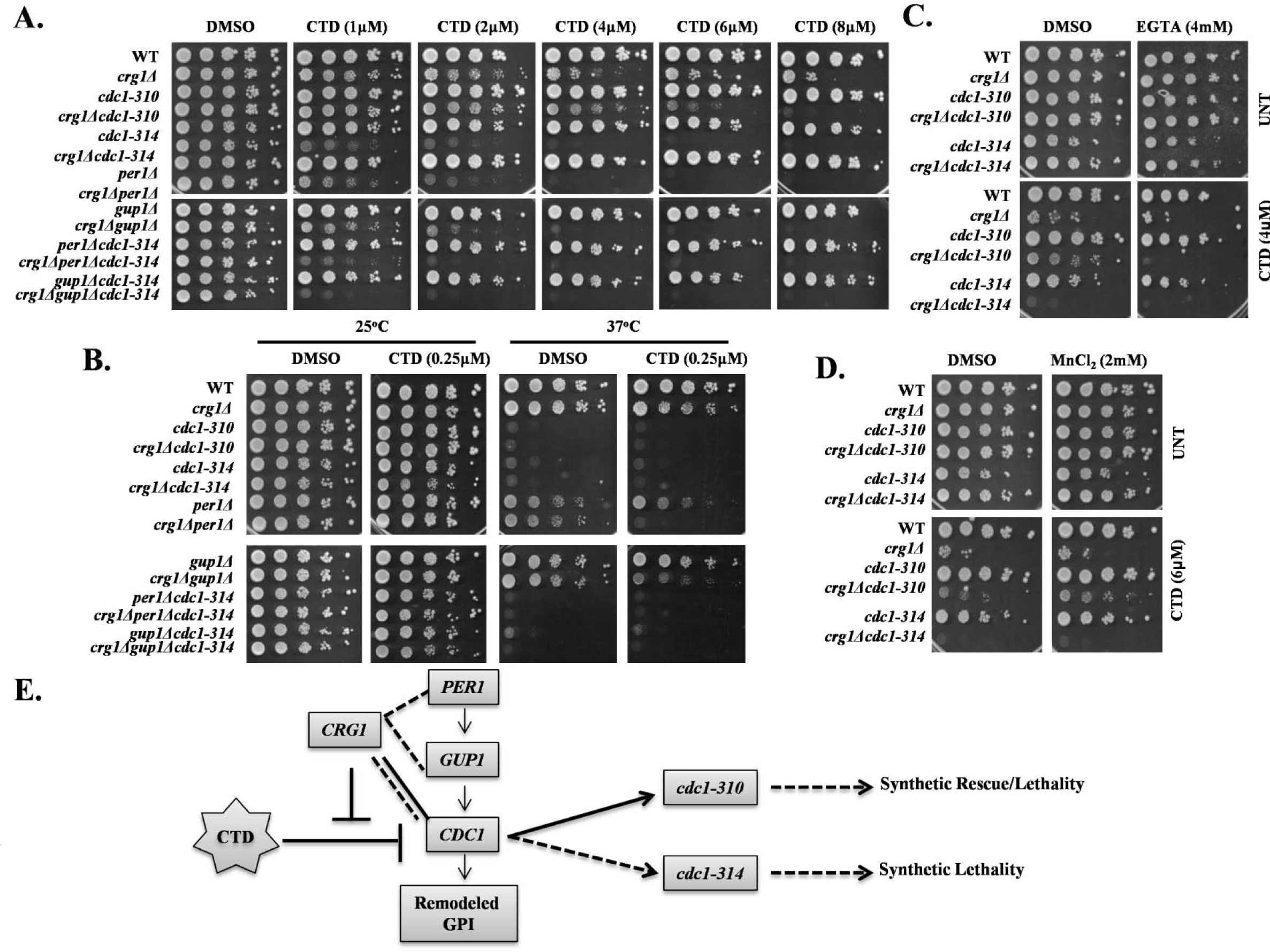
CTD targets Cdc1 activity involved in GPI-anchor remodeling. A, B, C, and D: Growth sensitivity assay. Equal numbers of serially diluted cells of WT and indicated mutants were spotted on SC agar media with various treatments. Images were captured after 72h of incubation. (A) *CRG1* shows synthetic lethality with GPI-anchor remodeling genes under CTD stress. Spot assay on media with increasing doses of CTD (1μM to 8μM) followed by incubation at 25°C. (B) *CRG1* shows synthetic lethality with GPI-anchor remodeling genes under CTD and heat stress. Spot assay on media containing CTD with or without ETA and the cells were incubated at 25°C and 37°C. (C) Mn^2+^ chelation increases CTD toxicity. Yeast strains indicated above were spotted on media containing CTD with and without EGTA and incubated at 25°C. (D) Mn^2+^supplementation decreases CTD toxicity. Yeast strains indicated above were spotted on media containing CTD and MnCl_2_ and incubated at 25°C. (E) Schematic representation of CTD dependent genetic interaction o *f CRG1* with GPI-anchor remodeling genes: *PER1, GUP1* and *CDC1.CRG1* show synthetic lethality with *PER1*, *GUP1* and *CDC1. cdc1-310* shows dose dependent interaction with *crg1Δ*; synthetic rescue at lower dose (2μM to 4μM) and synthetic lethality at higher dose (6μM to 8μM).

### CTD-induced phenotypes strongly correlate with loss of Cdc1 activity

Analysis of the phenotypes observed in this study and the investigations conducted previously suggest that CTD treatment induces phenotypes similar to *cdc1* mutants (*cdc1-314, cdc1-310*,* etc.*) (3,4,30). To get further evidence to support this, we employed genetics and cell biology-based experimental approaches. First, we measured the Slt2 phosphorylation upon CTD treatment. We observed that *cdc1-314* mutant showed increased phosphorylation of Slt2 compared to WT in untreated condition. On the other hand, CTD treatment induced the Slt2 phosphorylation in *crg1Δ, cdc1-314*, and *crg1Δcdc1-314* (Fig. 6A). Though CTD induced Slt2 phosphorylation in all the three mutants, the maximum level was measured in the double mutant *crg1Δcdc1-314*. We also observed decreased UPR in *cdc1-314* due to inhibition of *HAC1* mRNA splicing in untreated condition (Fig. 6B) (4). The splicing of *HAC1* mRNA decreased synergistically in the double mutant *crg1Δcdc1-314* with and without CTD treatment (Fig. 6B). Furthermore, we observed a defect in GPI-anchored protein sorting (Gas1-GFP) in *cdc1-314* (Fig. 6C) (3,4), which became worse if treated with CTD (Fig. 6C). Additionally, we found decreased expression of Gas1-GFP in *cdc1-314* (4). Interestingly, CTD treatment further decreased Gas1-GFP expression in *crg1Δ* and *crg1Δcdc1-314* (Fig. 6D and Fig. 4B). We found synergistic effect in all four phenotypes: Slt2 phosphorylation, UPR, Gas1-GFP localization, and Gas1-GFP expression, suggesting that Cdc1 might be a specific target of CTD.

**Figure 6:**
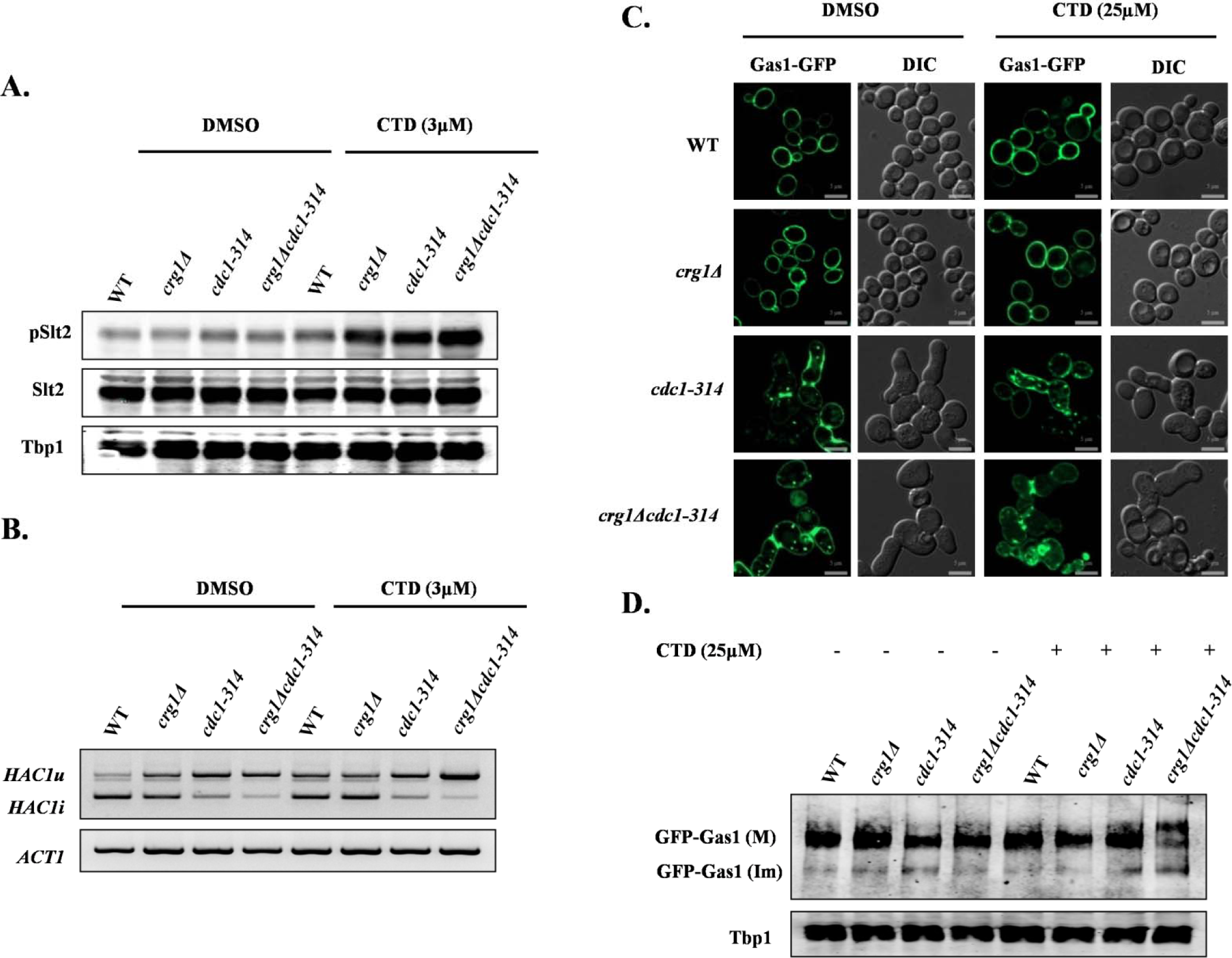
CTD treatment mimics *CDC1* mutation (*cdc1-314*). (A) CTD treatment induces Slt2 phosphorylation in *crg1Δ* and *cdc1-314* mutant. Western blot analysis of Slt2 phosphorylation in WT, *crg1Δ*, *cdc1-314* and *crg1Δcdc1-314* strains. Cells were grown at 25°C till mid-exponential phase, and then treated with CTD for 2h. (B) Synergistic inhibition of *HAC1* mRNA splicing in *crg1Δcdc1-314* mutant upon CTD treatment. RT-PCR analysis of *HAC1* mRNA in WT, *crg1Δ, cdc1-314* and *crg1Δcdc1-314* mutants. The cells were grown at 25°C till mid exponential phase and then treated with CTD for 2h. (C) CTD induces Gas1-GFP miss-sorting. Sub-cellular localization of Gas1-GFP in WT, *crg1Δ*, *cdc1-314* and *crg1Δcdc1-314.* Cells were transformed with YEp24-GAS1-GFP and grown in YPD media with and without CTD treatment for 6hr at 25°C. (D) CTD decreases expression level of Gas1-GFP. Western blot analysis of Gas1-GFP expression in cells described in (C).

Next, we measured the growth of GPI-anchor remodeling mutants in presence of CTD or anti-oxidants with increasing temperature. We found that the mutants of GPI remodeling genes (*per1Δ, gup1Δ, cdc1-310, cdc1-314, per1Δcdc1-314,* and *gup1Δcdc1-314*) were sensitive to a higher dose of CTD, and the sensitivity increased with increasing temperature (Fig. S7). We also found these mutants to be hypersensitive to reducing environment developed by supplementation of GSH or NAC into the medium (Fig. S7). The sensitivity to GSH as well as NAC increased again with elevated temperature. This result suggests that redox balance plays an essential role in remodeling process of the GPI-anchors. These results also provide an explanation for the synergistic lethal phenotype generated by the co-treatments with CTD+GSH or CTD+NAC (Fig. 2E). Additionally, ETA supplementation did not rescue the growth defect of *cdc1-314* and *cdc1-310* at higher temperature (Fig. S7), suggesting that ETA-mediated rescue in Gas1-GFP sorting in CTD-treated cells did not occur via the GPI-anchor remodeling mechanism. The hypersensitivity of the single mutant *cdc1-314* to the higher doses of CTD indicates the involvement of a *CRG1*-independent pathway targeted by the drug (Fig. S7). We also observed that a higher dose of CTD (300μM) altered Gas1-GFP sorting even in WT (Fig. S8A). Furthermore, to investigate whether CTD-induced alteration in GPI-anchored protein sorting was PP2A/PP1-dependent or not (24), we analyzed Gas1-GFP localization in *sit4Δ* (PP2A) and *GLC7/glc7Δ* (PP1) strains (Fig. S8B). We did not find any defect in Gas1-GFP localization in both the mutants, implying the CTD-induced alteration in GPI-anchored protein sorting was independent of PP2A and PP1. Over all, the CTD-induced phenotypes strongly correlate to that of *cdc1-314* allele, perhaps CTD targets Cdc1 activity.

### CTD alters GPI-anchored protein sorting in human cancer cells

The pathway for the biosynthesis and sorting of GPI-anchored proteins is conserved from yeast to higher eukaryotes (1,64). Therefore, we reasoned that the fundamental mechanism of action of CTD would be similar in yeast and human cells. To study the GPI-anchored protein sorting in human cells, we used GFP-CD59 as a model GPI-anchored protein (5). We observed that CTD induced aggregation of GFP-CD59 in HeLa cells, while the untreated cells showed normal distribution of the protein (Fig. 7A, S8C). This observation suggests that the molecular mechanism of action of CTD is conserved from yeast to human cells. Furthermore, we also checked the total expression of GFP-CD59 in HeLa cells upon treatment with CTD. Unlike yeast, we did not find any change in GFP-CD59 expression (Fig. 7B), suggesting a distinct mechanism for the clearance of aggregated proteins in human cells. Next, we measured the phosphorylation of p44/42 (a human homologue of yeast Slt2). We found a significant induction in p44/42 phosphorylation in HeLa and HepG2 cells upon CTD treatment (Fig. 7C). CTD treatment also decreased the *XBP1* mRNA expression (Fig. 7D), suggesting that the down regulation of UPR is similar to yeast cells (Fig. 2B, C) (65). To rescue the cells from CTD toxicity, we supplemented ETA, CHO and INO into the Dulbecco’s modified Eagles medium (DMEM). Interestingly, ETA supplementation rescued the HepG2 cells from CTD-induced cell death, but the rescue of HeLa cells was not significant (Fig. 7E, F), suggesting a cell-type-specific utilization of ETA due to different origin. On the other hand CHO and INO supplementation could not rescue the human cells (Fig. S11A,B,C,D) from CTD toxicity as observed in yeast cells (Fig 1, A). Since the phenotypes induced by CTD treatment in human and yeast cells are quite similar, we propose a conserved mechanism of action of CTD in eukaryotes.

**Figure 7:**
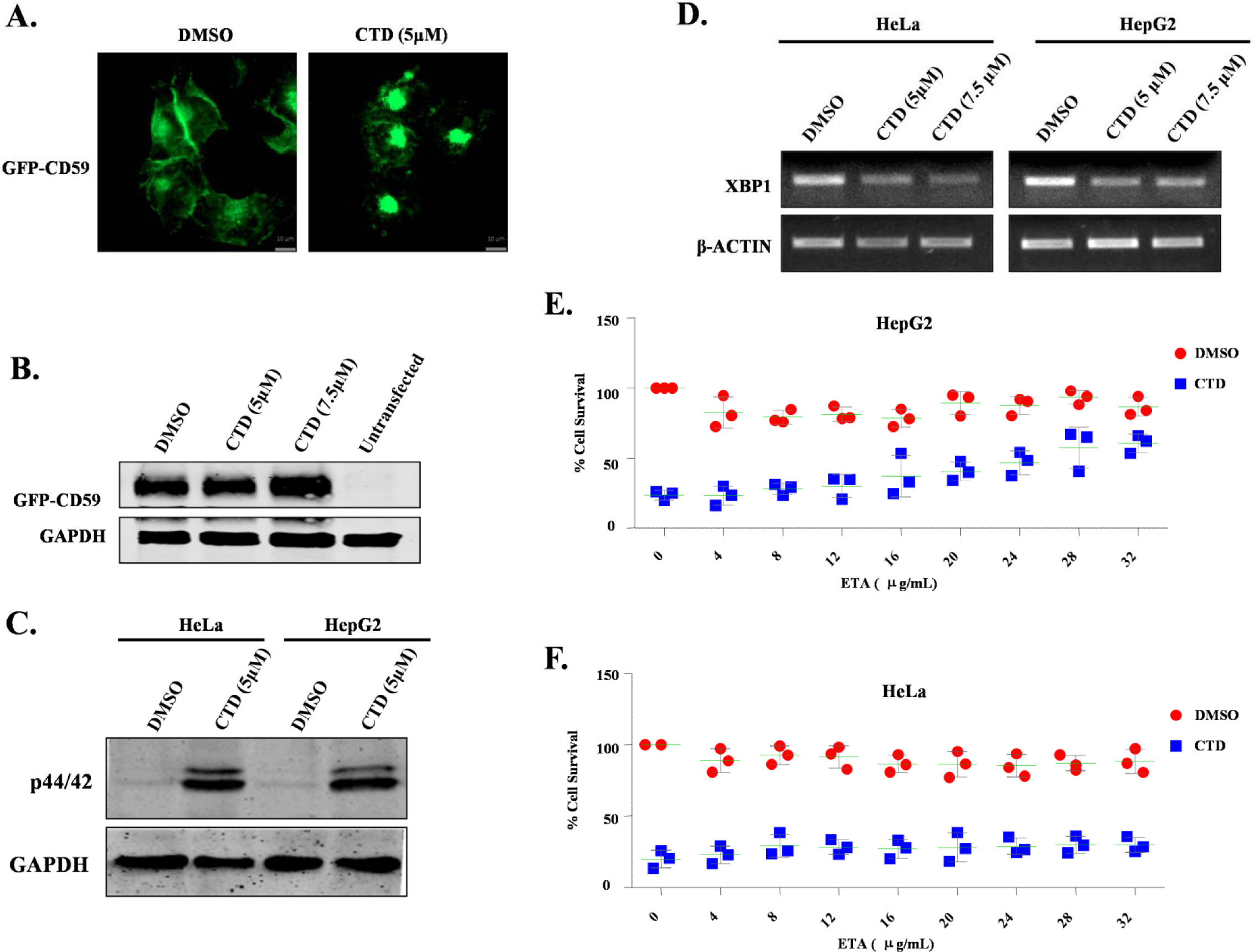
Conserved mechanism of CTD cytotoxicity in human cancer cells (HeLa and HepG2). (A) CTD alters GPI-anchored protein sorting. Microscopic visualization of GFP-CD59, stably expressing in HeLa cells with or without CTD treatment for 12h. (B) CD59 expression is unaffected with CTD treatment. Western blot analysis of GFP-CD59 expression in HeLa cells treated with CTD for 12h. (C) CTD treatment induces p44/42 (Slt2) phosphorylation. Western Blot analysis of p44/42 phosphorylation in HeLa and HepG2 cell lines after CTD treatment for 48h. (D) CTD treatment downregulates *XBP1* expression. Semi qRT-PCR analysis of *XBP1* expression in HeLa and HepG2 cell lines treated with CTD for 48h. (E) ETA supplementation rescues HepG2 cells from CTD cytotoxicity. MTT cell survival assay of HepG2 cells treated with CTD, supplemented with increasing concentrations of ETA. (F) ETA supplementation doesn’t rescue HeLa cells from CTD cytotoxicity. MTT cell survival assay of HeLa cells treated with CTD, supplemented with increasing concentrations of ETA.

**Figure 8:**
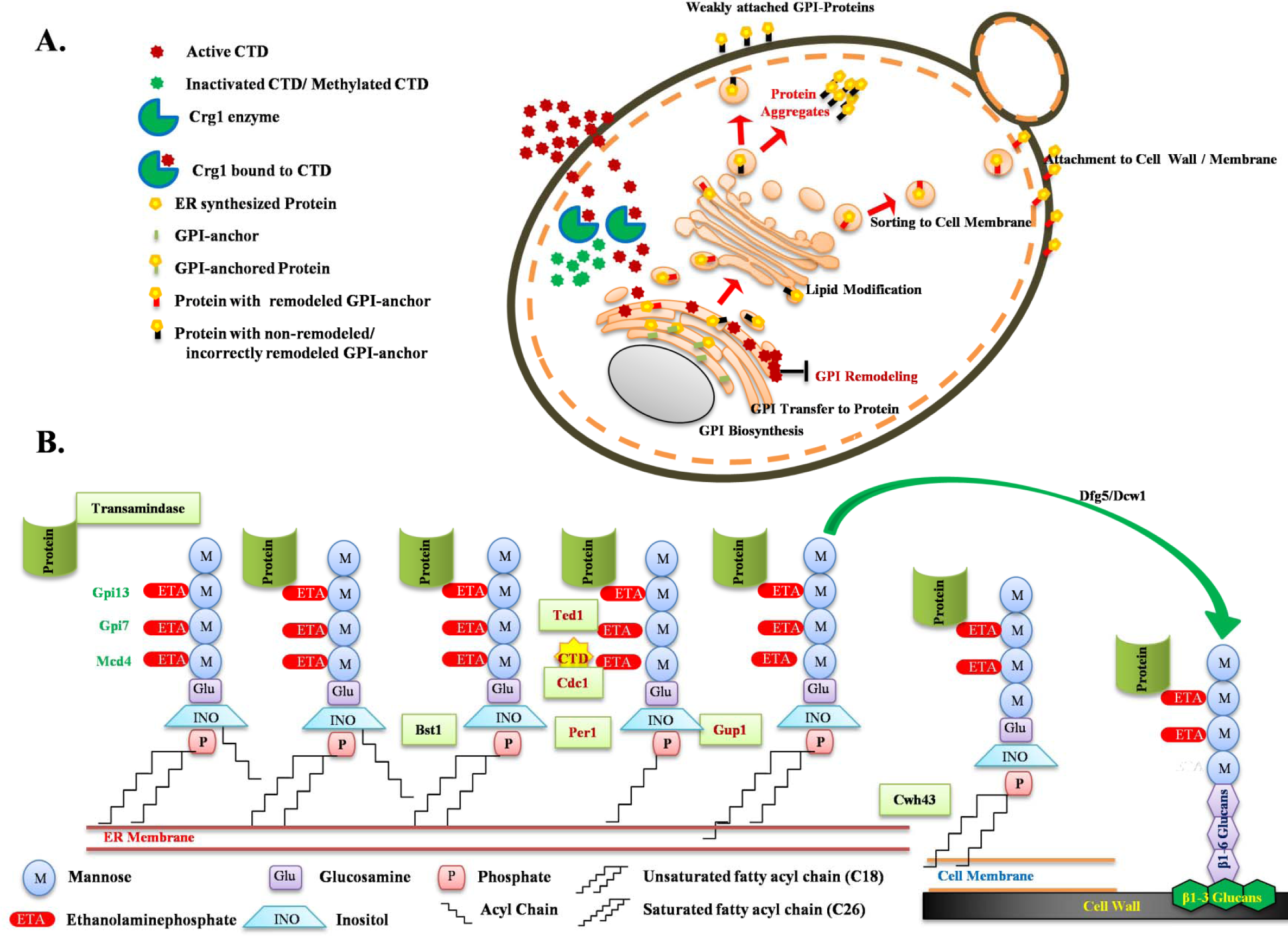
Schematic model illustrating the molecular targets and mechanism of CTD toxicity in yeast and higher eukaryotes. (A) The model describes yeast Crg1 as a key defense molecule, localized in the cytoplasm which protects the cell from CTD induced cytotoxicity by methyltransferase activity. Loss of Crg1 enhances the binding of CTD to its molecular targets and perturbs the related biological functions. In the absence of Crg1, CTD enters into the ER and disturbs the ER homeostasis by altering the GSH:GSSG ratio and GPI-anchor remodeling leading to missorting and aggregation of the proteins in the cytoplasm. (B) An illustration of the GPI anchor remodeling process in budding yeast. The C-terminal end of the protein is transferred to the ethanolaminephosphate of the third mannose of the GPI-anchor, catalyzed by a complex of enzymes GPI-transamidase. In the subsequent process Bst1 removes the acyl group from the inositol of GPI, Cdc1 removes ethanolaminephosphate from first mannose, Per1 removes the unsaturated fatty acid (C18:1) from the *sn-2* position of the GPI-lipid, Gup1 adds C26:0 saturated fatty acid at the *sn-2* position of the GPI-lipid and at last Cwh43 replaces the diacyglycerol type lipid with ceramide in GPI. Finally the GPI-anchor is transferred to plasma membrane or cell wall by Dfg5 or Dcw1. In this sequence of events CTD targets Cdc1 activity resulting into mislocalization and aggregation of GPI anchored proteins.

## DISCUSSION

GPI-anchored proteins control essential biological functions in animal cells by regulating the cell–to-cell communication, adhesion, and signal transduction (2,9). GPI-anchored proteins are also shown to be involved in tumorigenesis and metastasis (10-12). Targeting an essential cellular pathway is one of the key aspects of anticancer chemotherapeutics. In this study, employing extensive genetic and cell biological approaches, we identified Cdc1 (yeast homologue of human PGAP5)-mediated GPI-anchor remodeling as a mechanistic target of CTD in addition to PP2A and PP1. Biochemical validations will further support our observations.

CTD has been shown to disturb phospholipid homeostasis in *crg1Δ* mutant (30). To understand the underlying mechanism of its action, we screened the *crg1Δ* mutant for the auxotrophy of different phospholipids upon CTD treatment. This approach helped us to conclude that CTD specifically affects PE, which can be rescued by exogenous supplementation of ETA. CTD treatment induced phenotypes similar to *psd1Δ* (39,44), and we found that *PSD1* was synthetically lethal in combination with*CRG1* under CTD stress. The reason of PE auxotrophy upon CTD treatment may be either inhibition of PE biosynthesis or alteration in PE-associated structures e.g. GPI-anchors. The biosynthesis of PC mainly depends on the availability of PE in ER, which suggests that PE deficiency can lead to the deficiency of PC as well (66). However, the supplementation with PC did not rescue the growth defect induced upon CTD treatment. Thus, we conclude that CTD probably alters the PE-associated structures or functions rather than PE biosynthesis. PE deficiency is also known to induce ER stress and UPR in yeast (44). Contrarily, we found decreased UPR upon CTD treatment in the absence as well as presence of the UPR inducers (TM and DTT). Our further investigations revealed that the drop in UPR upon CTD treatment was due to alteration in ER-redox homeostasis and Cdc1 activity (4) where we found that increased GSH level or lack of Cdc1 activity diminished UPR. The oxidative environment in ER is maintained by low GSH:GSSG (1:1 to 3:1) ratio for correct folding and modifications of the proteins (47,49,50). Our study demonstrates that the oxidative environment is also essential for the process of GPI-anchor remodeling (Fig. S7). ER is the site of synthesis and fate determination of the secretory proteins in the cell. Biosynthesis and maintenance of the yeast cell wall majorly depends on these secretory proteins (1). We believe that the CTD-induced cell wall damage (30,34) is due to alteration in ER homeostasis. The synergistic lethal effect of CTD with ER stress inducers (Heat, DTT, and TM) and cell wall perturbing agents (CR and CFW) support this hypothesis. CTD-induced Slt2 phosphorylation also increases synergistically with ER stress inducers. Thus, we conclude that CTD-induced ER stress triggers cell wall damage. We also found rescue from CTD-induced cell wall damage upon ETA supplementation, the reason of which is the restoration of the GPI-anchored protein sorting (38,44). However, we do not know the exact mechanism by which the increased PE level restores the GPI-anchored protein sorting against CTD.

Next, we investigated the molecular mechanism for the ER stress and cell wall damage upon CTD treatment. The genetic interaction profile of *CRG1* suggests that the ER-Golgi traffic system is a major pathway affected by CTD (30). In yeast cells, the proteins that travel from the ER to cell wall are mostly the GPI-anchored proteins. GPI-anchored proteins constitute a major part of the total cell wall proteins and are required for the biosynthesis and maintenance of the yeast cell wall (1,2,7,58,64). Alteration in biosynthesis or sorting of GPI-anchored proteins induce ER stress and cell wall damage (4). Interestingly, we observed mis-sorting and aggregation of the GPI-anchored protein (Gas1-GFP) upon CTD treatment. *CRG1* showed synthetic rescue with GPI-anchor biosynthesis genes (*GPI2*, *GPI13*, and *MCD4*) and synthetic lethality with GPI-anchor remodeling genes (*GUP1*, *PER1*,and *CDC1*) upon CTD stress, indicating that the CTD alters GPI-anchored protein sorting by targeting the remodeling process (4). These results also support the genetic interaction profile of *CRG1* reported previously (30). Besides, we identified *CDC1* as an additional new gene that showed synthetic lethality with *CRG1* in the presence of CTD. *CDC1* encodes for Mn^+2^-dependent mannose-EtNP phosphodiesterase required for the removal of EtNPfrom the first mannose of the GPI-anchor (4). The *crg1Δcdc1-314* double mutant shows strong sensitivity to CTD compared to *crg1Δ* and *cdc1-314* single mutants. The triple mutant strains*crg1Δper1Δcdc1-314* and *crg1Δgup1Δcdc1-314* were found to be even more sensitive to CTD compared to single (*crg1Δ, per1Δ*, *gup1Δ, cdc1-310*, and *cdc1-314*) and double mutants (*crg1Δper1Δ* and *crg1Δgup1Δ*), suggesting that GPI-anchor remodeling is the major target of CTD. On the contrary, another allele of *CDC1*, *cdc1-310*, shows a dynamic phenotype upon CTD treatment. It shows synthetic rescue at lower dose and synthetic lethality at higher dose of CTD. Such dynamic and contrasting phenotypes of the two different alleles of *CDC1* suggest a possibility of direct interaction of the enzyme with the small molecule CTD. To get more evidence in support of this hypothesis, we manipulated the Mn^+2^concentrations in the medium. We found that CTD toxicity enhanced with decreasing concentrations of Mn^+2^ in the medium and vice versa, indicating an essential requirement of the Cdc1 activity to tolerate the CTD toxicity. Based on these results, we believe that CTD inhibits Cdc1 activity. CTD shows stronger affinity to Cdc1-314 than Cdc1-310, probably due to the specific protein confirmation. Previous studies suggest that CTD acts as a potent inhibitor of protein phosphatasesPP2A and PP1 (23,24,67). However, our observations suggest that it can also inhibit lipid phosphatases such as Cdc1. CTD-dependent synthetic lethality of *SAC1* (phosphatidylinositol phosphate phosphatase) with *CRG1* supports this hypothesis (30,68). Furthermore, the *sit4Δ* (PP2A) and *GLC7/glc7Δ* (PP1) mutants do not show any defect in GPI-anchored protein sorting, suggesting that CTD-induced alteration in GPI-anchored protein sorting is independent of its known protein targets PP2A and PP1 (23,24,30). Furthermore, we also found that the higher dose of CTD induces the same phenotypes in WT and *cdc1-314* as it does in *crg1Δ* mutant at sub-lethal dose, suggesting that CTD targeted pathways are independent of *CRG1*.

The enzymes involved in GPI-anchor biosynthesis and remodeling in yeast are mostly conserved in higher eukaryotes, suggesting that CTD can act through a similar mechanism in higher eukaryotes. To validate the existence of a conserved mechanism of CTD toxicity, we extended our studies to human cell lines HeLa and HepG2. We observed similar phenotypes induced by CTD in human cells. We found missorting and aggregation of GPI-anchored GFP-CD59 in the cytoplasm of HeLa cells upon CTD treatment, which was very similar to that of Gas1-GFP in yeast. Similarly, CTD also induced phosphorylation of p44/42 (yeast Slt2), supporting the previous observations of CTD-mediated activation of different MAPKs (18,21). We also found decreased expression of XBP1 (yeast *HAC1*) upon CTD treatment which might be via ATF6 signaling that regulates the target gene XBP1 (65). The similar phenotypes produced by CTD in yeast and human cell lines suggest that the drug functions through a conserved mechanism.

Our study provides explanations to various observations reported upon CTD treatment in different organisms. CTD-induced alteration in GPI-anchored protein sorting can be a reason for the acantholysis (69-71) and inhibition of cancer metastasis (12). CTD-induced perturbation in adhesion, morphogenesis, and membrane trafficking in *Candida albicans* may be due to alteration in GPI-anchored protein sorting (72). The G2/M cell cycle arrest by CTD (18,19,67) is probably due to inhibition of Cdc1 activity as the loss of Cdc1 functions also induces G2/M cell cycle arrest (73). CTD has been a traditional medicine to cure warts and molluscum contagiosum caused by viral infections. Our study suggests that CTD can be further explored as an antiviral or antiprotozoan drug, utilizing its property of altering ER-Golgi traffic system (8,15,74).

Since CTD targets a conserved and essential pathway, its exposure can also lead to lethal side effects. Therefore, the drug delivery is required to be very specific. A cancer- or tumor-specific delivery of CTD is the only way to make it a successful chemotherapeutic anticancer drug. Similarly, the poisoning of cattle foods by the contamination with the blister beetle is another big challenge, as there is no antidote available against the beetle toxin. Our study suggests ETA can serve as a potent antidote against CTD poisoning. In summary, we identified a novel target of CTD in addition to PP2A/PP1 and a potent antidote that neutralizes its lethal toxic effect.

## MATERIALS AND METHODS

### Yeast strains, plasmids and growth conditions

Unless otherwise stated, *Saccharomyces cerevisiae* strains, used in this study were isogenic with S288c (BY4741 or BY4743). All the strains, plasmids and primers used in this work are listed in Tables: S1-S3 respectively. Yeast strains were grown in Synthetic Complete (SC) or Yeast Peptone Dextrose (YPD) media at 30°C, maintaining the optimum growth excluding some temporal stress conditions. Various reagents used in different experiments were purchased from Sigma, Merck, Himedia, Invitrogen, BioRad and Applied Biosystems.

### Growth sensitivity assays

**Serial dilution assay:** Equal number (OD_600_ = 1.0) of overnight grown yeast cells were serially diluted, 10 fold for 5 times, and then spotted on SC agar media. The spotted cells were incubated at different temperatures according to the various experimental conditions.

**Growth curve assay:** Equal number (OD_600_ = 0.2) of exponentially growing yeast cells were inoculated in 96-well plates with and without different treatments and grown for 23-28h in the automated plate reader (Biotek) acquiring reading at OD_600_ in an interval of every 30 minutes.

### Preparation of yeast whole cell protein extract for western blot analysis

Protein extraction from yeast cells was done by following the TCA protein extraction protocol (75). The equal number of cells were harvested and washed twice with 20% of TCA. Cell pellets were re-suspended in 20% of TCA with an equal volume of glass beads and vortexed rigorously to lyse the cells. TCA precipitated protein extract was washed with ethanol, resuspended in 0.5M of Tris-Cl (pH 7.5) with 1X loading buffer. The sample was boiled at 100°C for 10 minutes and centrifuged at maximum for 10 minutes to remove the debris. The supernatant was taken ahead for western blot analysis. The primary antibodies used in this study for immunoblotting experiments were: Anti-phospho-p44/42 (Cell Signaling, Catalog 4370), Anti-Mpk1 (Santa Cruz Biotechnology Inc., Catalog SC-6803), Anti-GFP (Sigma, Catalog G1544), Anti-GAPDH (Cell Signaling, Catalog 5174S). Primary antibody used against Tbp1 was polyclonal antisera raised in rabbit.

### Cell surface protein extraction

Yeast cells were washed twice with sodium phosphate buffer (0.1 M, pH 8.0). Wash-out solution was kept at 4°C. Collected cells were re-suspended again in sodium phosphate buffer with 2mM of DTT and incubated at 4°C for 2h maintaining gentle agitation. Now the cells were pelleted down by centrifugation, and the supernatant along with the washout fraction was precipitated using 20% TCA in the final volume. TCA precipitated cell surface proteins were separated via 8%- SDS-PAGE and stained with 0.1 % CBBR (63).

### RNA extraction, cDNA synthesis and qPCR for *HAC1* mRNA splicing

RNA isolation was performed by using the heat/freeze RNA isolation protocol (76). Briefly; cells were grown until mid-exponential phase, harvested by centrifugation and washed twice with 1xPBS. Harvested cells were lysed with 1% of SDS in AE buffer (50mM Sodium acetate, 10mM EDTA, pH 5.3). An equal volume of acidic phenol of pH 4.2 was added, incubated at 65°C for 4 minutes followed by freezing at −85°C for 4 minutes, centrifuged for 2 minutes at maximum speed to separate the aqueous layer. The aqueous phase was mixed with equal volume of phenol:chloroform:isoamyl alcohol (PCI; 25:24:1) and separated again from the phenol phase. The total RNA present in aqueous phase was precipitated by adding sodium acetate (0.3M) and 2.5 volume of absolute ethanol. The cDNA synthesis was done by following the standard protocol provided by iScript™ cDNA Synthesis Kit (BioRad, Catalog 1708891). *HAC1* mRNA splicing was measured using primer specified in Table S3, following the PCR conditions as described previously (53).

### β-galactosidase assay

Exponentially growing yeast cells were harvested and washed twice with Lac-Z buffer (10mM KCl, 1mM MgSO4, 50mM β-mercaptoethanol, and 100mM NaPO4, pH 7.0). Cells were lysed using 0.01% SDS, and 22.7% chloroform in Lac-Z buffer in final volume of 250μl. Subsequently 500 μl of ONPG (2mg/ml) was added and incubated at 30°C till the appearance of pale yellow color. The reaction was quenched by adding 500μl of sodium bicarbonate (1M). The reaction mixture was centrifuged at maximum speed for 15 minutes, and the supernatant was collected to measure the absorbance at the wavelength of 420nm. Miller unit for the β-galactosidase activity was determined by applying the formula: Miller Unit = [OD_420_/OD_600_*Time (Min)]*1000 (45).

### Fluorescence microscopy

Fluorescence microscopy was done to study the sorting of GPI-anchored protein (Gas1-GFP) with or without CTD. Yeast cells were grown in YPD medium for different time points and harvested by centrifugation (6000rpm, 2 minutes, 4°C). Cell pellets were resuspended in PBS and kept on ice at least for 30 minutes (59). Gas1-GFP localization was observed using the ZEISS-Apotome.2 fluorescence microscope under 60x oil emulsion objective lens. For the microscopic localization study of the EGFP-FCD59, the overnight grown HeLa cells with 50% confluency were treated with CTD (5μM) for 12 hours and visualized under 20x emulsion oil objective lens (5).

### Cell culture and maintenance of Human cell lines

Human cell lines (HeLa and HepG2) were maintained in Dulbecco’s modified Eagle’s medium (DMEM, Lonza) having 10% fetal bovine serum (FBS, Gibco) and antibiotics,*i.e.,* penicillin (100U/ml) and streptomycin (100μg/ml). Both the cell lines were grown at 37°C with 5% CO_2_.

### Cell survival assay

Percentage survivability of the cells against CTD exposure was measured by MTT assay. HeLa and HepG2 cells were seeded in 96 well plate equal in number (5000 cells) in each well. Cells were incubated for 24 h. Media was removed, and fresh media was added to the cells, simultaneously cells were also challenged with CTD with or without supplementation of ETA for 48 h. 10μl of MTT solution (5mg/ml in 1xPBS) was added and incubated at growth conditions for four hour. 100μl of DMSO was added and mixed well. Absorbance was recorded at 570nm using microplate reader (Biotek) (16).

### Statistical data analysis

Statistical analysis for all the data of this study was performed by using GraphPad Prism-5 software. Each graph shows the individual data points with mean value as a horizontal green line. The error bars represents mean±SD of minimum three individual repeats. We applied Two-way ANOVA, Bonferroni post test where *= p<0.05, **= p<0.01 and ***= p<0.001 and ns=p>0.05.

## ACKNOWLEDGEMENTS

We thank Benjamin S. Glick for providing us the mutants of GPI-anchor remodeling pathway (*cdc1-310, cdc1-314, per1Δ, gup1Δ, per1Δcdc1-314, gup1Δcdc1-314* and WT-JK9-3d). We also thank Vishal M. Gohila for gifting us the *psd1Δ* and WT (BY4741). We are also thankful to Peter Walter, Laura Popolo and Morihisa Fujita for gifting us pPW344 (UPRE-LacZ), YEp24-GAS1-GFP and pME-NeodHmEGFP-F-CD59_hGHss respectively. We acknowledge all the members of the lab for critical suggestions and helpful discussions throughout the work. DBT-India supported the JRF/SRF fellowship to PKS. The work was supported for funding from IISER Bhopal and SERB Govt. of India to RST.

## AUTHOR CONTRIBUTIONS

The hypothesis and experiments were designed by RST and PKS. Experiments were performed by PKS and the results were analyzed by PKS and RST. Manuscript was written by PKS and RST.

